# Handy divisions: Hand-specific specialization of prehensile control in bimanual tasks

**DOI:** 10.1101/2024.08.28.610133

**Authors:** Anvesh Naik, Satyajit Ambike

## Abstract

When hammering a nail, why do right-handers wield the hammer in the right hand? The dynamic dominance theory suggests a somewhat surprising answer. The two hands are specialized for different types of tasks: the dominant for manipulating objects, and the non-dominant for stabilizing objects. Right-handers wield the moving object with their right hand to leverage the skills of both hands. Functional specialization in hand use is often illustrated using examples of object manipulation. However, the dynamic dominance theory is supported by wrist kinematics rather than object manipulation data. Therefore, our goal was to determine whether this theory extends to object manipulation.

We hypothesized that hand-specific differences will be evident in the kinematics of hand-held objects and in the control of grip forces. Right-handed individuals held two instrumented objects that were coupled by a spring. They moved one object while stabilizing the other object in various bimanual tasks. They performed motions of varying difficulty by tracking predictable or unpredictable targets. The two hands switched roles (stabilization vs movement) in various experimental blocks. The changing spring length perturbed both objects. We quantified the movement performance by measuring the objects’ positions, and grip force control by measuring grip-load coupling in the moving hand and mean grip force in the stabilizing hand.

The right hand produced more accurate object movement, along with stronger grip-load coupling, indicating superior predictive control of the right hand. In contrast, the left hand stabilized the object better and exerted a higher grip force, indicating superior impedance control of the left hand. Task difficulty had a weak effect on grip-load coupling during object movement and no effect on mean grip force during object stabilization. These results suggest that dynamic dominance extends to object manipulation, though the weak effect of task difficulty on grip characteristics warrants further investigation.

## Introduction

Approximately 90% of humans favor their right hand for manual tasks [1]. This asymmetry in hand use has intrigued scientists since the time of Woodworth [2], and has led to the development of two major theories explaining the motor control behind handedness. The global dominance theory posits that the dominant (right) hand is better in all manual activities compared to the non-dominant (left) hand; and because each hand is controlled mostly by the contralateral hemisphere, this theory suggests overall superiority of the left hemisphere in right-handed individuals for controlling both hands [3-6]. In contrast, the dynamic dominance theory proposes that each hemisphere specializes in different aspects of movement control [7-9]. The left hemisphere specializes in predictive control; it anticipates changes in the environment and arm dynamics and generates coordinated movements with the right arm better than the right hemisphere can produce coordinated movements with the left arm. In contrast, the right hemisphere specializes in impedance control; it stabilizes the left arm against perturbations better than the left hemisphere can stabilize the right arm. This view was based on seminal behavioral data gathered about four decades ago [10, 11], which Sainburg and colleagues subsequently formalized into the dynamic dominance theory based on wrist kinematics during reaching movements in healthy individuals [8, 9, 12] and unilateral stroke survivors [13, 14].

Here, our primary goal was to move beyond wrist kinematics to determine whether the dynamic dominance theory explains the manipulation of hand-held objects. More importantly, we were interested in determining whether the control of the forces exerted by the digits on the object was also described by this theory. It is plausible that hemispheric specialization for controlling arm movements extends to the control of digit force, but this is not inevitable for two reasons. First, non-overlapping neural substrates are involved in the control of arm movement and digit forces [15-20]. Second, behavioral evidence indicates that the control of arm movements and grip forces is partly dissociated [21]. For example, humans adjust the grip force on a static object *before* they start moving the object, indicating that grip forces can change even when the object’s motion, and therefore arm motion, is the same [22]. Therefore, behavioral evidence for dynamic dominance in object manipulation is essential for establishing the domain of applicability of this theory.

Our strategy to look for this evidence was to inspect the grip force-load force coupling (henceforth grip-load coupling for brevity) and the magnitude of grip force. These variables are critical for manipulating objects stably, and they reflect the predictive and impedance control aspects of prehension. The predictive aspect of the control is demonstrated by humans coupling the grip force with the predicted loads on the grasped object [23-28]. The impedance aspect is demonstrated by humans exerting grip force exceeding the minimum required to ensure a slip-free grasp [25, 27, 28]. A higher grip force also stiffens the wrist through muscle co-contraction, further improving the object’s positional stability [28-31]. Thus, if dynamic dominance extended to grip force control, then the grip-load coupling would be stronger in the dominant (right) hand when moving an object and the grip force magnitude would be higher in non-dominant (left) hand when stabilizing an object.

Additionally, our secondary goal was to quantify how task difficulty interacts with dynamic dominance during bimanual prehensile tasks. Recent studies on unimanual prehension reported that grip force characteristics are altered by task difficulty. Grip-load coupling is stronger when individuals track unpredictable compared to regular and predictable paths with a handheld object [32, 33]. Furthermore, the grip force increases when external loads act on the object, but it increases more when the loads are variable [34]. We argue below that dynamic dominance may not manifest for simple tasks; therefore, we systematically studied how dynamic dominance is influenced by task difficulty.

Previous studies on grip force control lacked evidence for dynamic dominance, possibly due to task design issues. Jaric and colleagues used isometric bimanual force production tasks with fixed handles [35-39]. They found similar grip-load coupling in both hands, and lower grip force in the left hand [38], results that are incompatible with the predictions of dynamic dominance theory [8, 40]. The issue here may have been that the static tasks in this study favored the left hand and masked effects that could emerge while moving objects.

Other studies involved object movement, but still failed to observe between-hand differences. Participants exhibited similar grip-load coupling and grip force in both hands while moving two objects with different weights [41, 42] or holding one object steady while simultaneously moving another object [43, 44]. These results may stem from ceiling effects; the tasks may have been too simple, and both hands could perform the tasks equally well without significantly changing grip forces [45].

These studies revealed two crucial considerations for studying dynamic dominance in prehension. First, tasks should include both movement and stabilization to leverage the specialization of each hand. Second, the task must be sufficiently challenging to reveal hand-specific differences in grip force control. Based on this thinking, we designed a bimanual task that approximated bread slicing: one hand stabilized an object against the disturbances arising from the movements of the other hand [12]. Participants manipulated two objects, one per hand, connected by a spring. They tracked a visual target by moving one object while holding the other object stationary. The target profiles were predictable or unpredictable in different conditions, which altered task difficulty. This task challenged the motor system by requiring distinct actions from each hand while accounting for the spring-induced disturbances.

Our working hypothesis was that dynamic dominance would extend to object kinematics and grip force control. The right hand will show superior object movement accuracy because the left hemisphere specializes in predictive control (hypothesis H1). In contrast, the left hand will show superior object stabilization because the right hemisphere specializes in impedance control (hypothesis H2). These hypotheses extend the classical observations on wrist kinematics [8, 12] to the control of hand-held objects. Furthermore, the grip-load coupling strength – the predictive aspect of prehension – will be higher for the right hand when moving the object (hypothesis H3). The mean grip force – the impedance aspect of prehension – will be higher for the left hand while stabilizing the object (hypothesis H4). Finally, these grip force characteristics will show an interaction between hand and task difficulty. A more difficult task will strengthen the grip-load coupling in both hands while moving objects, but the increase will be greater in the right hand than the left (hypothesis H5). In contrast, a more difficult task will increase the mean grip force in both hands while stabilizing objects, but the increase will be greater in the left hand than the right (hypothesis H6).

## Materials and Methods

### Participants

Twenty-four healthy, young individuals (11 females; age = 24.7 ± 4.6 years; weight = 73.2 ± 14.6 kg; height = 168.7 ± 10.6 cm) volunteered to participate in the study. Handedness was assessed by the Edinburgh Inventory [46]. This inventory provides a laterality quotient (LQ) ranging from -100 (strong left-handedness) to +100 (strong right-handedness). All participants were right-hand dominant (LQ = 88.5 ± 9.5), with no previous history of neuropathies or trauma to the upper limbs. All participants provided informed consent in accordance with the procedures approved by the Institutional Review Board of Purdue University.

### Equipment

Participants held two instrumented objects in a pinch grasp with the tips of index finger and thumb of each hand (Fig 1). Six-component force transducers (Nano 17-E, ATI Industrial Automation, Garner, NC) mounted on the objects measured the force of each digit. The distance between the grasping surfaces of the two force transducers on an object was 6.5 cm. Sandpaper (100C medium grit) was glued to the surface of transducers to increase the coefficient of friction between the transducer and the digits. At the start of each experimental session, force transducers were zeroed with objects resting vertically on the table, and no digits contacting the sensors. A tension spring (stiffness 19.5 N/m, and resting length 8 cm) was attached between the two objects (Fig 1). When the spring was at its resting length, the horizontal distance between the centers of the sensors on the two objects was 13 cm. The total weight of the apparatus was 2 N. Four reflective markers were attached to each object (Fig 1) and a four-camera motion capture system (Vicon Vero VE22-S, Oxford, UK) was used to track the position of both objects. The motion capture system was calibrated for each participant to obtain a tracking error less than 1 mm inside the capture volume. The MotionMonitor software (Innovative Sports Training Inc.) was used to synchronize and collect output signals from the force transducers (sampled at 1000 Hz) and the motion capture system (sampled at 200 Hz).

**Fig 1.**
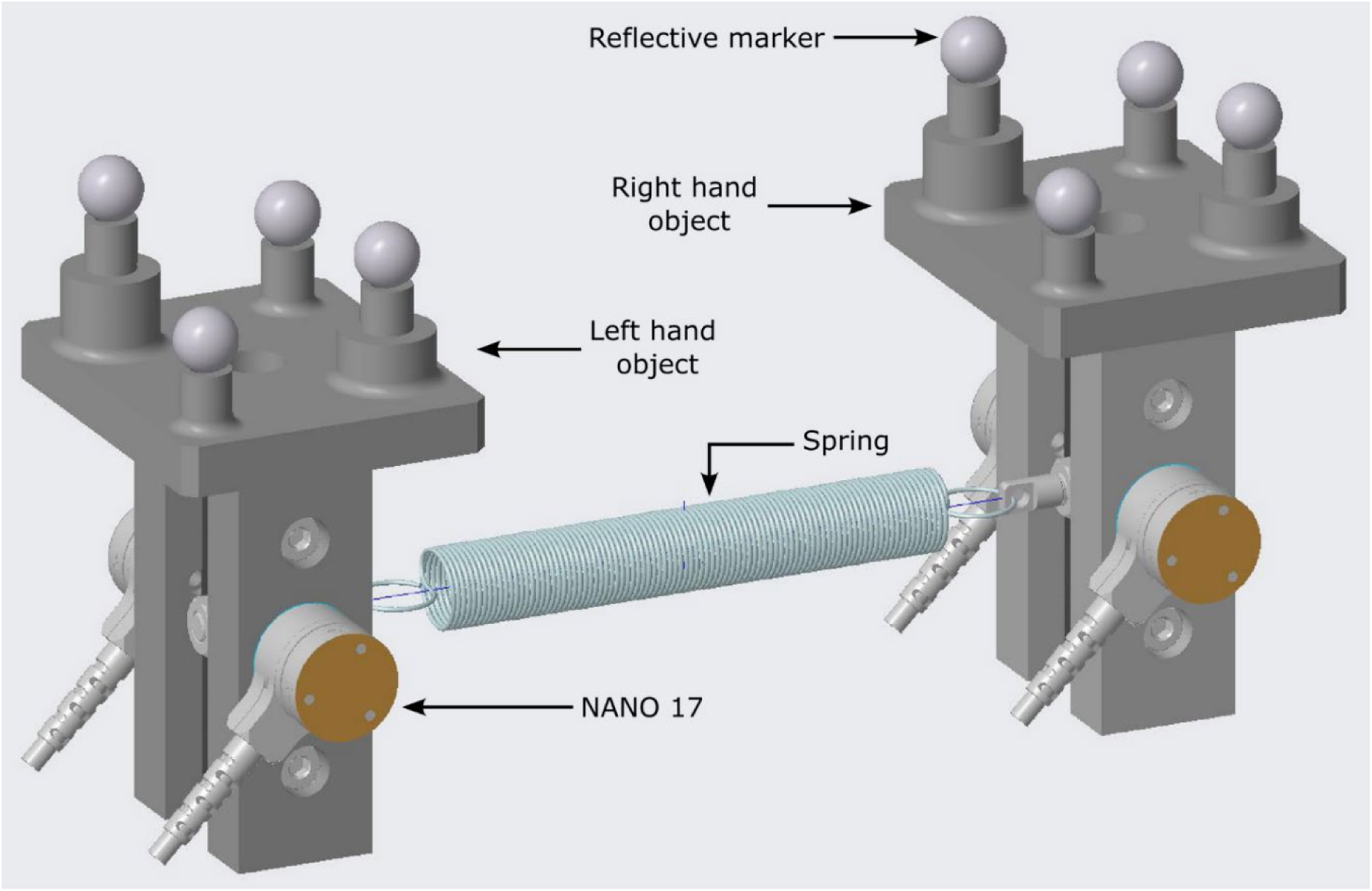
Illustration of the instrumented objects. Nano 17 force sensors recorded digit forces. A motion capture system tracked the positions of four reflective markers fixed to each object.

### Experimental setup and procedure

Before the start of an experimental session, participants cleaned the digit tips of both hands using alcohol wipes to normalize the skin condition. Participants sat upright on a piano bench facing a computer screen placed on a table (Fig 2). A reference point was marked on the table as the origin for the world co-ordinate system. The positive X axis pointed to the participant’s right, the positive Y axis pointed along the anterior direction, and positive Z axis pointed vertically upward.

**Fig 2.**
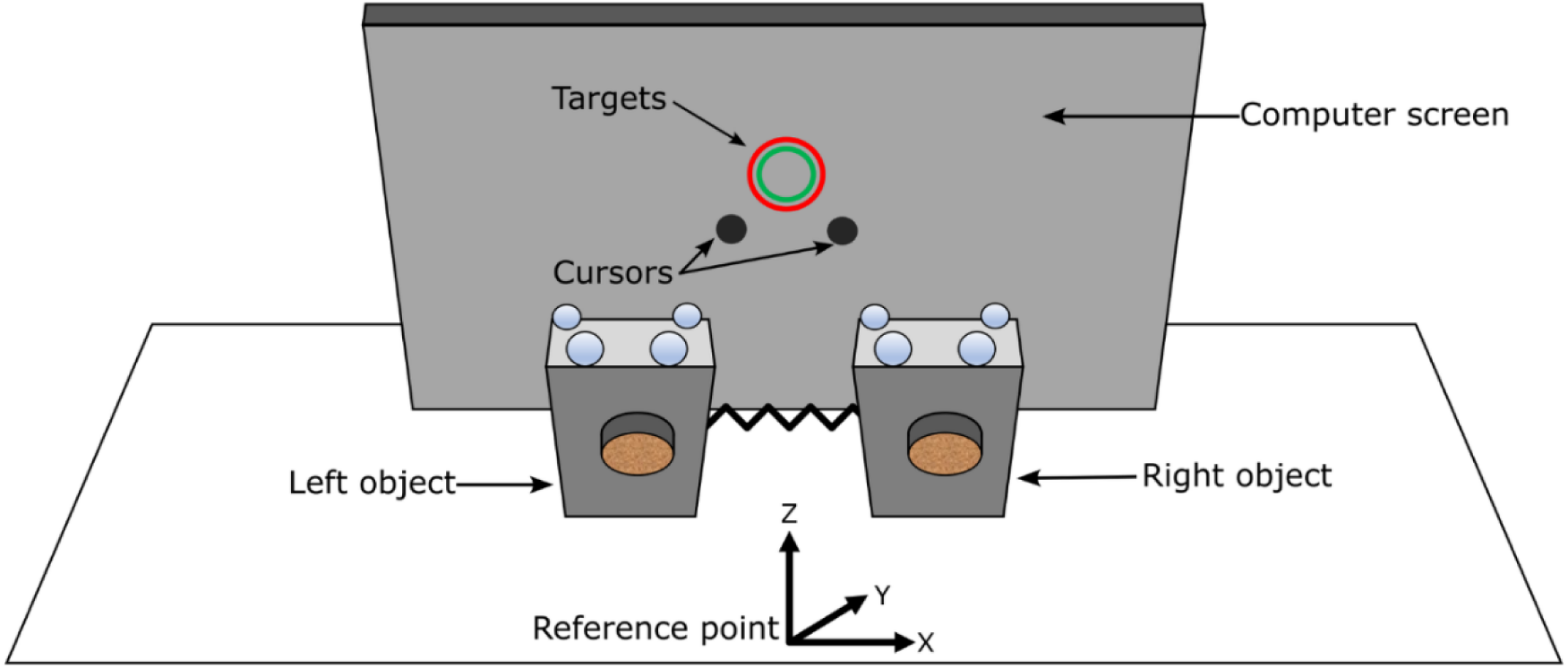
Experimental setup. Participants held the objects in pinch grasps and positioned them above the reference point. Targets and cursors were displayed on the computer screen placed on the table.

The participants grasped both objects by placing the index finger and the thumb of each hand on the transducers. They lifted the objects and held them vertical and stationary at a comfortable height above the reference point (Figs 2 and 3A). The computer screen displayed two hollow circular *targets* and two solid circular *cursors* (Fig 3A). The cursors indicated the real-time positions of the objects in the frontal X-Z plane. The centroid of the four markers on the object was defined as the position of that object. The cursors were flipped on the screen so that the cursor on the left represented the right-hand object and vice versa. Although the initial positions of cursors were flipped, their motions on the screen were veridical. That is, the rightward motion of any of the objects moved the corresponding cursor to the right on the screen, and so on for the other horizontal and vertical directions (Fig 3A). Participants moved both objects vertically and apart from one another until the two cursors overlapped at the center of the targets (Fig 3B). In this position, the spring was stretched by approximately 3 cm which generated a baseline horizontal spring force of approximately 0.6 N on each object.

**Fig 3.**
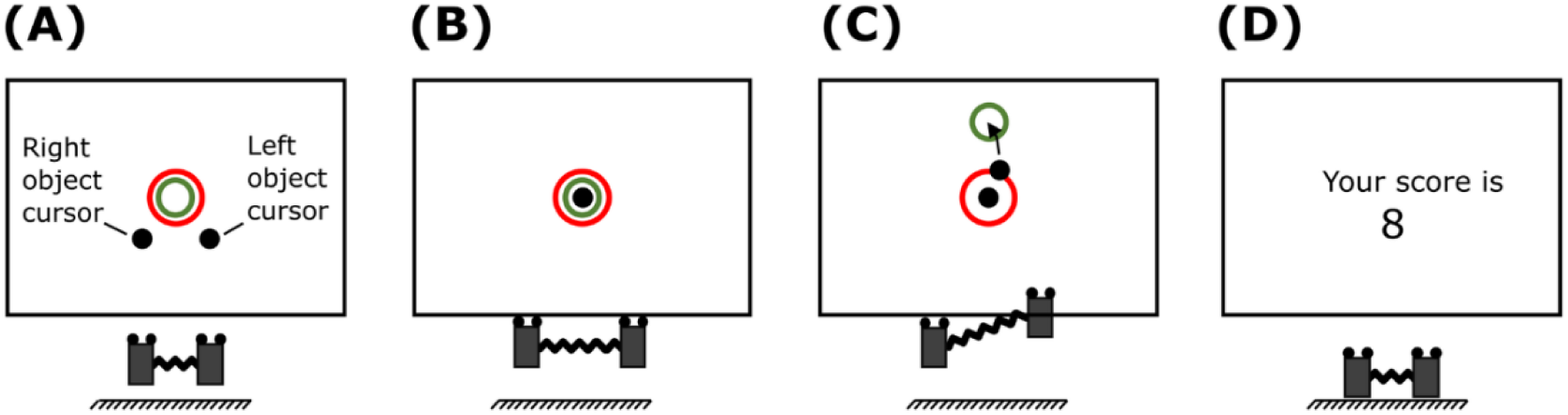
Targets and cursors feedback on the computer screen. (A) position of cursors and objects at the start of a trial, (B) objects positioned in the frontal plane such that the spring is stretched to bring both cursors within the targets, (C) target tracking with right object while left object is stabilized, and (D) score displayed at the end of the trial.

In each trial, the green target moved vertically on the screen, whereas the red target remained stationary (Fig 3C). Participants were instructed to move one object vertically so that the corresponding cursor tracked the green target as accurately as possible, while simultaneously stabilizing the other cursor inside the stationary red target (Fig 3C). The red target was bigger in size than the green target to prevent both targets from overlapping. At the end of each trial, targets and cursors vanished and participants replaced both the objects on the table and released the grasp.

There were four blocked experimental conditions (2 *hand* × 2 *task difficulty*), with repeated trials in each block. In two blocks, the right hand moved one object, and the left hand stabilized the other object. In the other two blocks, the role of each hand was switched. The task difficulty was modulated by changing the moving target’s trajectory. In different blocks, the movement profile was a regular and predictable triangular wave (amplitude 0.24 m, frequency 0.5 Hz). In the other two blocks, the profile was an irregular and unpredictable fractional Brownian motion (fBm; Hurst exponent 0.25). The two profiles and the tracking performance by each hand by one representative participant are shown in Fig 4. The order of each *hand* × *task difficulty* block was counterbalanced across participants using the Latin square design.

**Fig 4.**
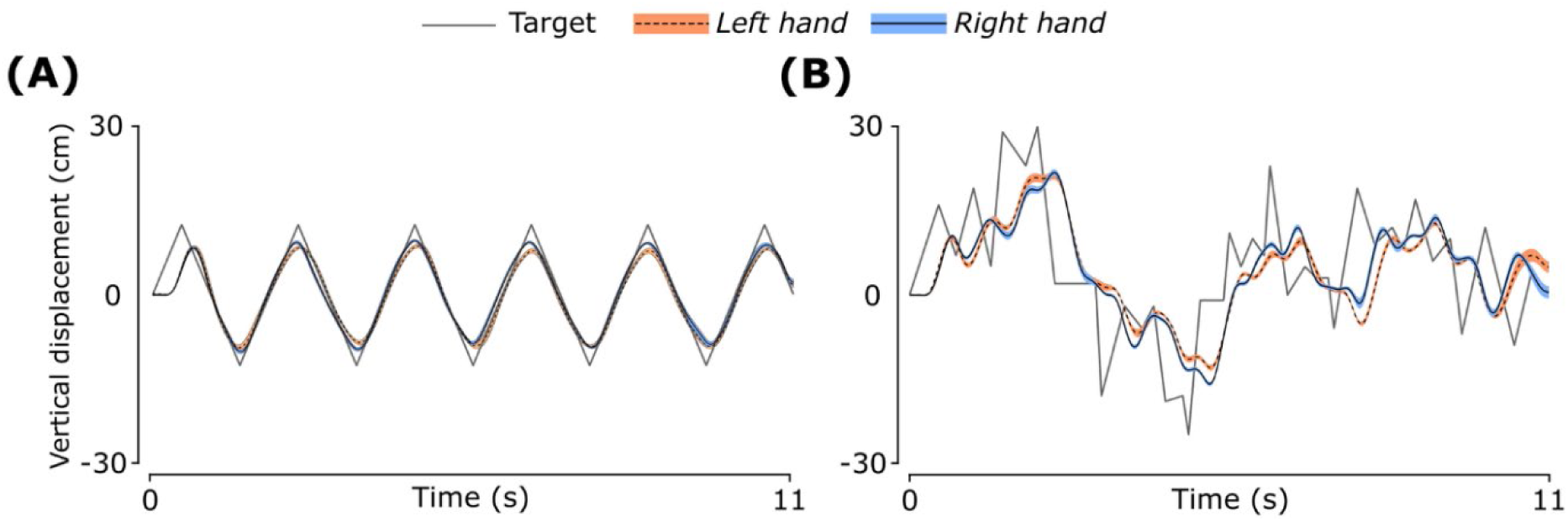
Target profiles and tracking performance by each hand. (A) Regular (triangular) movement and (B) Irregular (fBm) movement. Data shown are across trial mean and standard error for a representative participant.

There were 10 trials in each block and 40 trials in total during the experimental session. Each trial was 15 s long. In the first 4 seconds, the targets remained in the initial position and participants stabilized both cursors inside the targets (Fig 3B). The target movement began at the 4^th^ second and lasted until the end of the trial. At the start of each block, participants were informed which hand to move and which hand to stabilize. Participants were given adequate practice before each block till the experimenter was satisfied with the task performance. After each trial, a score reflecting the participant’s performance was displayed on the screen (Fig 3D). The score was the sum of the mean squared error between the center of each cursor to the center of the corresponding target. Participants were instructed to keep this score as low as possible.

Participants rested for 15 s between trials, and for 60 s between blocks. Additional rest periods were provided when participants felt fatigue in their arms or digits, and they were encouraged to ask for additional rest whenever required. The entire protocol lasted for about an hour. None of the participants requested additional rest, and none of the participants reported fatigue during the protocol.

### Data analysis

Custom MATLAB programs were written for data analyses (R2023a, The MathWorks Inc). To match the sampling frequencies of the kinematics and the digit forces, the force data was downsampled to 200 Hz. The down-sampled finger forces and kinematic data were low-pass filtered at a cutoff frequency of 4 Hz using a fourth-order, zero-lag Butterworth filter [33]. The data within an *analysis window* between 4 s and 15 s was used for further analysis.

The grip force for each hand was computed as the sum of the index finger and thumb normal forces of that hand [47]. The load force on each object was computed as the vector sum of tangential forces along the sensor-digit interface on both force transducers on an object. The inertial force vector was computed for the moving object as the object’s mass times its acceleration in the X-Z plane. The spring force on the moving object was then obtained by subtracting the load and weight vectors from the inertial force vector. Since the spring connected the two objects, the same spring force acted on the stabilized object as well. Profiles of the grip and load forces from a representative participant are shown in Fig 5.

**Fig 5.**
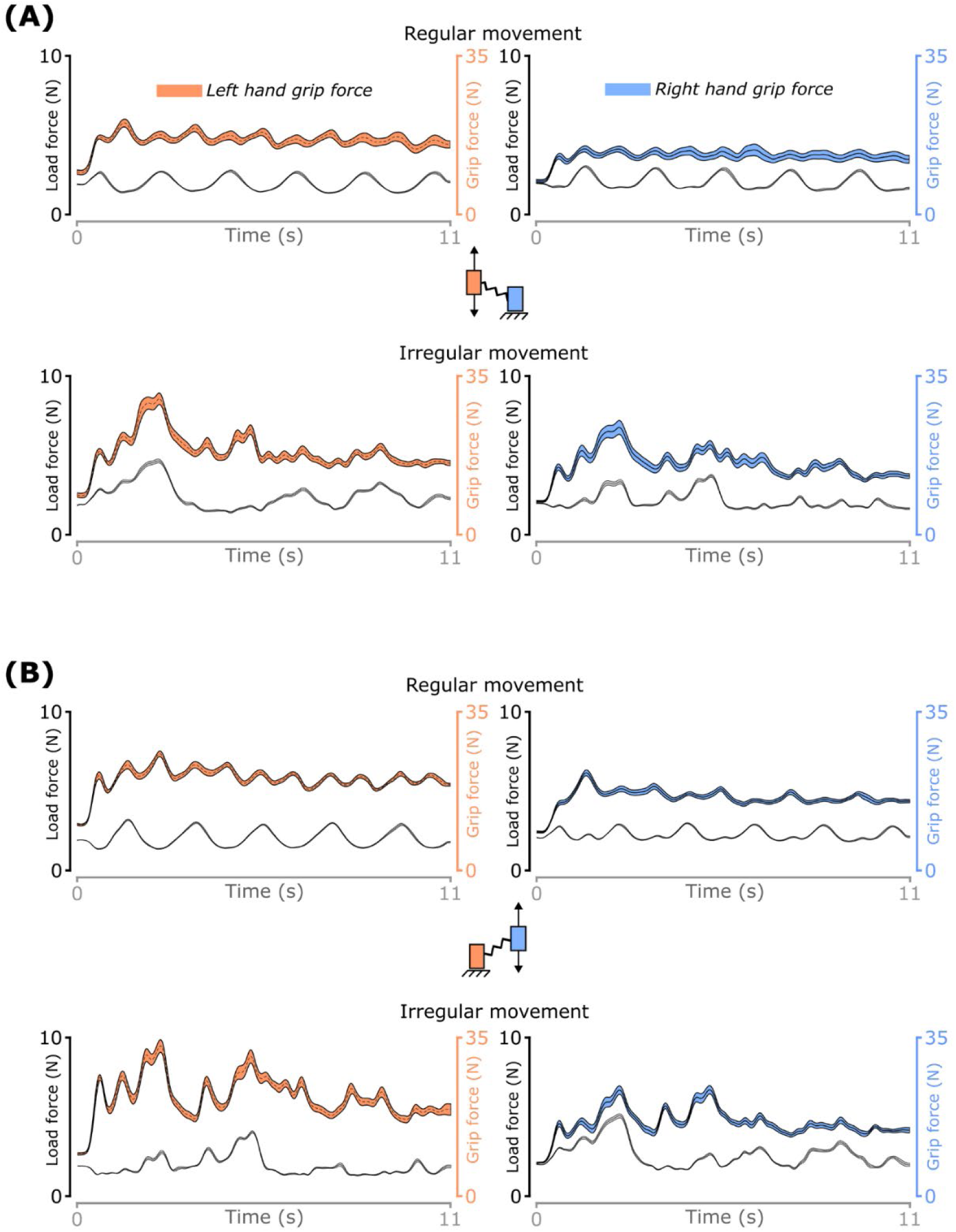
Grip and load forces for both hands. (A) moving left and static right hand during regular and irregular movements with the left hand, (B) moving right and static left hand during regular and irregular movements with the right hand. Data shown are across trial mean and standard error for a representative participant.

#### Basic performance measures

To quantify *task difficulty*, we computed performance score for each trial. To test whether participants stretched the spring before movement initiation, the spring length at the start of the analysis window (4 s after trial initiation) was quantified. To determine whether the orientation of both objects was vertical, we quantified object tilt as the angle between the force transducer’s plane and the X-Y plane since only this tilt influenced the load force. Finally, we computed the mean and variance of the load force on both objects within the analysis window to quantify the effect of *task difficulty* on characteristics of load force.

#### Task performance measures

To assess tracking performance of the moving hand, the root mean squared error (RMSE) between the centers of the moving target and the cursor were computed separately along the vertical and mediolateral directions. A lower RMSE along the vertical indicates better tracking performance and lower RMSE along the mediolateral direction implies straighter movements.

To assess the performance of the stabilizing hand at maintaining the object’s position, the standard deviation of the object’s position was computed in the frontal plane. A difference in the standard deviation across conditions could arise from difference in the external load. Therefore, the standard deviation was normalized by the spring force over the trial to obtain the compliance of the stabilized object [12]. The spring force was computed as the mean of the magnitudes of the spring force vectors within a trial. Technically, the external load on the object is the sum of the spring force and the object weight. As the object weight was constant, it was not included in the compliance calculation. Lower standard deviation and compliance imply better stabilization.

#### Measures of grip force control

The grip-load coupling was quantified for the moving hand using linear cross-correlational analysis and a non-linear technique called cross-recurrence quantification analysis (CRQA). The mean grip force was computed for the stabilizing hand by averaging the grip force over the analysis window for each trial.

##### Cross-correlational analysis

Linear cross-correlational analysis was performed to determine the correlation at zero lag [48]. We computed correlation at zero lag and not maximum correlation because approximately 80% trials (772 out of 960 trials) had zero lag between grip and load forces.

##### Cross-recurrence quantification analysis

Although cross-correlational analysis has been effective in capturing gross features of the coupling [23, 48], these methods are not suited to identify subtle temporal variations in the coupling [32, 33, 49]. Therefore, we also used a non-linear method called CRQA to quantify the strength of the grip-load coupling.

CRQA quantifies how two processes unfold and interact over time by computing how often their time series come close to each other in a common reconstructed phase space [50]. This technique proceeds in three steps. First, a common phase space for both time series is constructed using time-delayed versions of both series [51]. Second, instances when the phase-space trajectories of the two time series are within a pre-defined distance, known as radius, from each other are identified. These instances, called cross recurrence points, are used to create a cross recurrence plot. Third, the patterns in the cross recurrence plots are characterized by computing the appropriate outcome measures.

To construct the phase space and the cross recurrence plot for the digit forces, the maximum (L_∞_) norm [51] was used. The three input parameters required for this technique were identified using established recommendations: the embedding dimension [52], time delay [53], and radius [54]. These parameters were computed using all trials across all participants. The embedding dimension was set to 5, and the time delay was set to 23 samples (115 ms). The augmented Dickey-Fuller test [55, 56] identified non-stationarity in the grip force time series. Therefore, the radius was selected adaptively for each trial to ensure a fixed recurrence rate of 5% [33, 57].

We consistently observed vertical line structures in the cross-recurrence plots (Fig 6). This suggest that the state of one process does not change or changes slowly relative to the state of the other process, i.e., one time series gets trapped at a location while the other deviates from that location [58]. This implies the coupling between the two processes is intermittent [32, 59], and the degree of intermittency can be used to quantify the strength of the coupling. In particular, the strength of the coupling is inversely proportional to the intermittency, which, in turn, can be quantified using characteristic length of the vertical lines. This characteristic length was computed using trapping time, which is the average length of all the vertical lines in a plot [59, 60].

**Fig 6.**
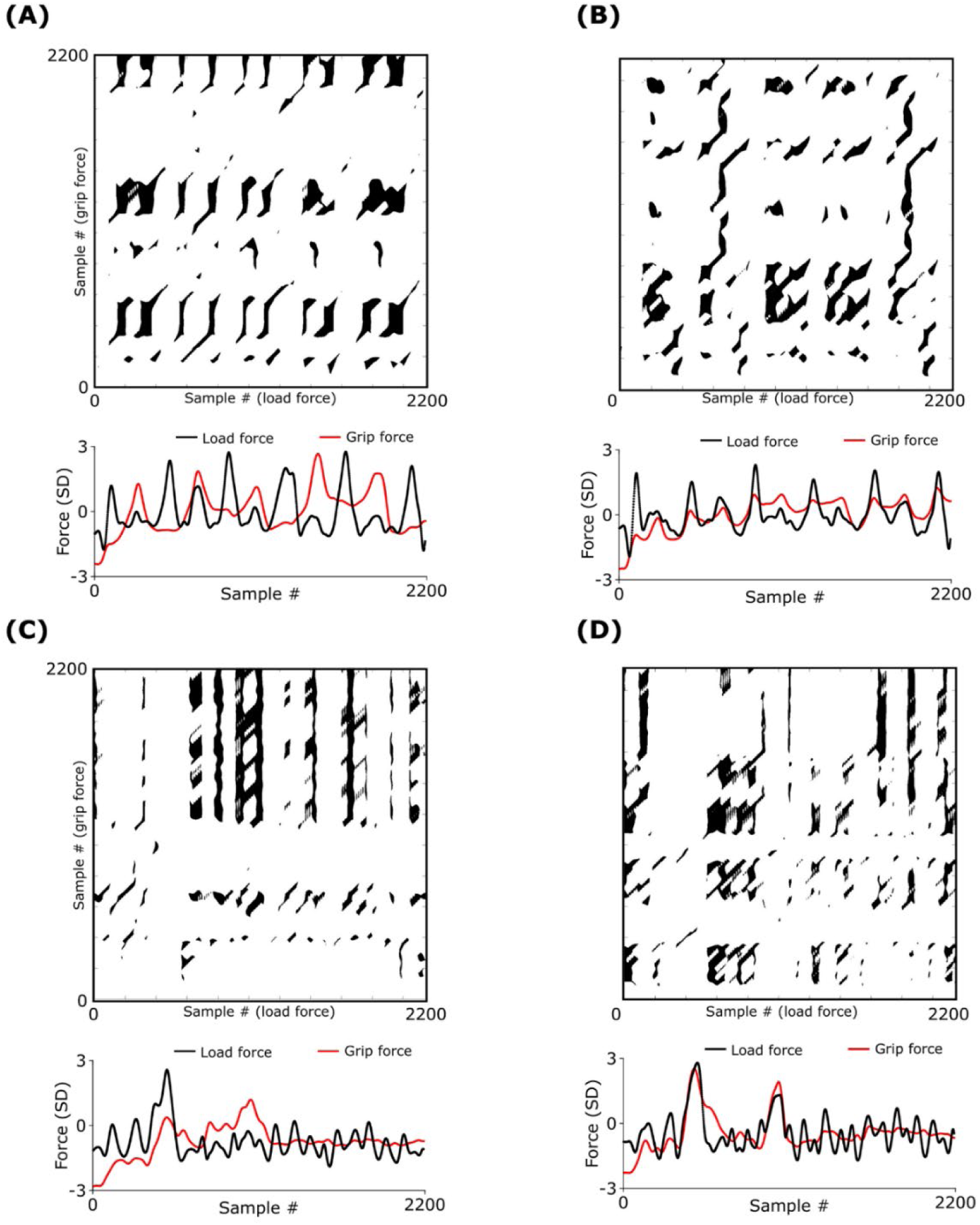
Cross-recurrence plots for a representative subject. (A) Regular movement with left hand, (B) regular movement with right hand, (C) irregular movement with left hand, and (D) irregular movement with right hand. Normalized grip and load forces are shown below each cross recurrence plot.

### Statistics

The data are presented as mean ± standard errors (SE) in the Results section, unless mentioned otherwise. All outcome measures were averaged across 10 trials for each *hand × task difficulty* block. Normality and constant variance requirements were checked visually by plotting studentized residual plots for each averaged outcome measure. The Box-Cox transformation was applied when the requirements were violated. The correlation coefficient values were z-transformed to meet requirements of normality. However, non-transformed data are presented in the Results section. Then, separate linear mixed-effects (LME) models [61] were fit to each basic performance measures, task performance measures, and measures of grip force control. The two experimental factors (*hand × task difficulty*) and their two-way interactions were included as fixed effects and *participant* was included as a random effect. Tukey-Kramer test was used to perform post-hoc pairwise comparisons when significant interaction effects were observed. Effect sizes were quantified by computing Cohen’s *d*. All statistics were performed using the SAS statistical software (version 9.4; SAS Institute, Cary, NC), with an α-level of 0.05.

## Results

### Basic performance measures

Participant score (Table 1) was not affected by *hand* (F_(1,23)_ = 0.7; *p* = 0.4) but was affected by *task difficulty* (F_(1,23)_ = 4182; *p* < 0.01). Post-hoc analysis revealed that the score was lower during regular movement compared to irregular movement (Cohen’s d = 13).

**Table 1.** Summary of participant performance scores for each *hand* × *task difficulty* block. The values computed are across participant average ± S.E. within each block.

|  | Left – regular | Right – regular | Left – irregular | Right – irregular |
| --- | --- | --- | --- | --- |
| Mean ± S.E.<br>(cm<sup>2</sup>) | 8.43 ± 0.5 | 7.81 ± 0.4 | 77.6 ± 1.9 | 78.7 ± 2.3 |

The stretched spring length before movement initiation was not affected by *hand* (F_(1,23)_ = 1.7; *p* = 0.2) or *task difficulty* (F_(1,23)_ = 0.3; *p* = 0.60), when both objects were stabilized with the respective cursors inside the green target (see Fig 5B in Methods). The center-to-center distance between the objects before movement initiation averaged across *hand* and *task difficulty* and participant was 16 cm ± 0.2 cm, indicating that participants stretched the spring initially by 3 cm as required.

The mean tilt of the moving object (Fig 7A) was affected by *hand* (F_(1,23)_ = 17.0; *p* < 0.01), but not by *task difficulty* (F_(1,23)_ = 0.21; *p* = 0.65). Post-hoc analysis revealed that the mean tilt was lower for the right object compared to the left object (Cohen’s *d* = 0.8). The mean tilt of the stabilized object (Fig 7B) was also affected by *hand* (F_(1,23)_ = 20.5; *p* < 0.01) but not by *task difficulty* (F_(1,23)_ = 0.39; *p* = 0.53). Post-hoc analysis revealed that the mean tilt was lower for the right object compared to the left object (Cohen’s *d* = 0.9).

**Fig 7.**
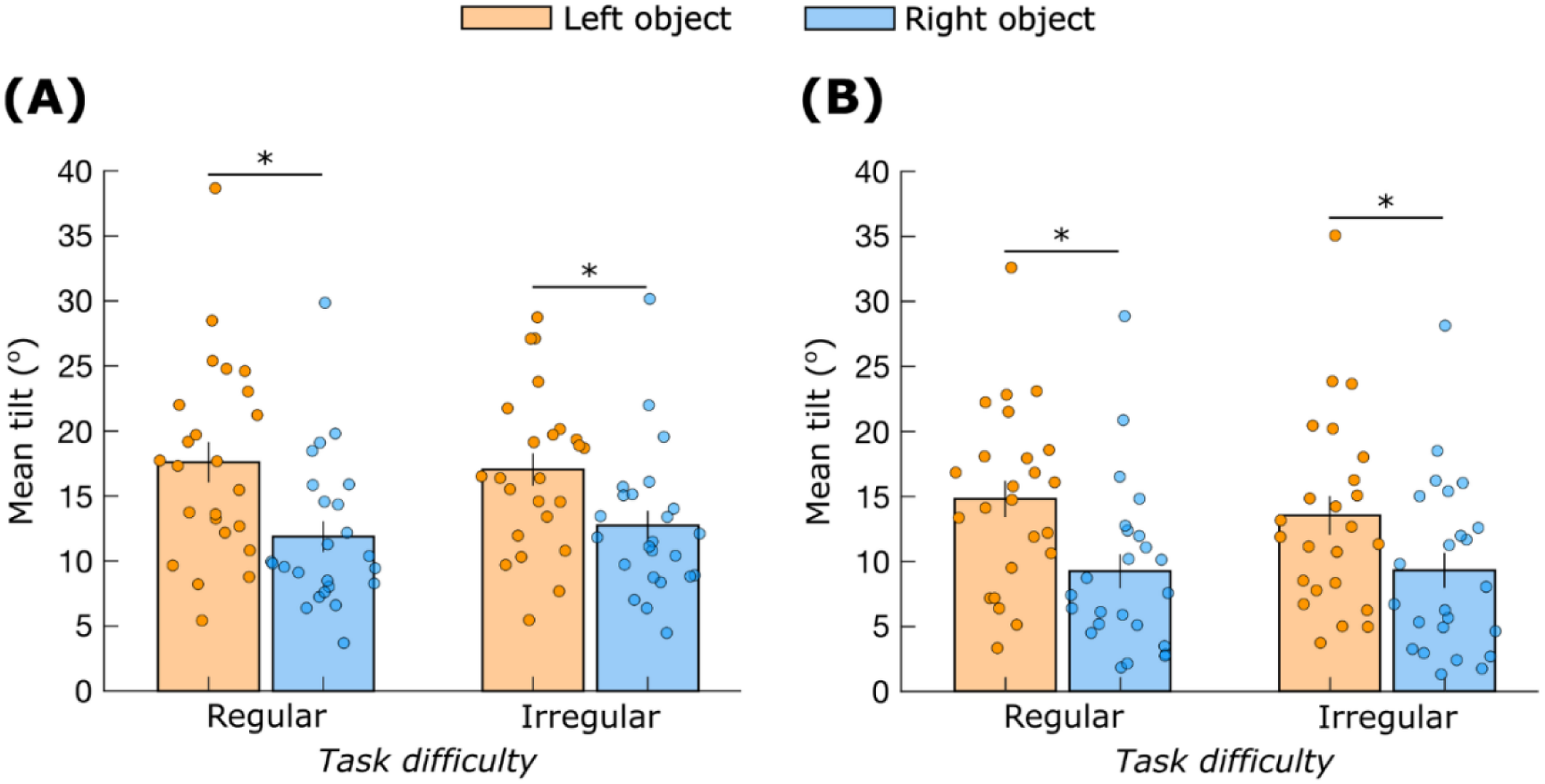
Mean object tilt. (A) Moving hand, and (B) Static hand. ‘*’ indicates significant differences (*p* < 0.05). Data are mean ± standard error.

The mean load force on the moving object (Fig 8A) was affected by *hand* (F_(1,23)_ = 4.9; *p* = 0.04) and *task difficulty* (F_(1,23)_ = 74.0; *p* < 0.01). Post-hoc analysis revealed that the mean load force was higher for the right object compared to the left (Cohen’s *d* = 0.4) and lower during regular movement compared to irregular movement (Cohen’s *d* = 1.7).

**Fig 8.**
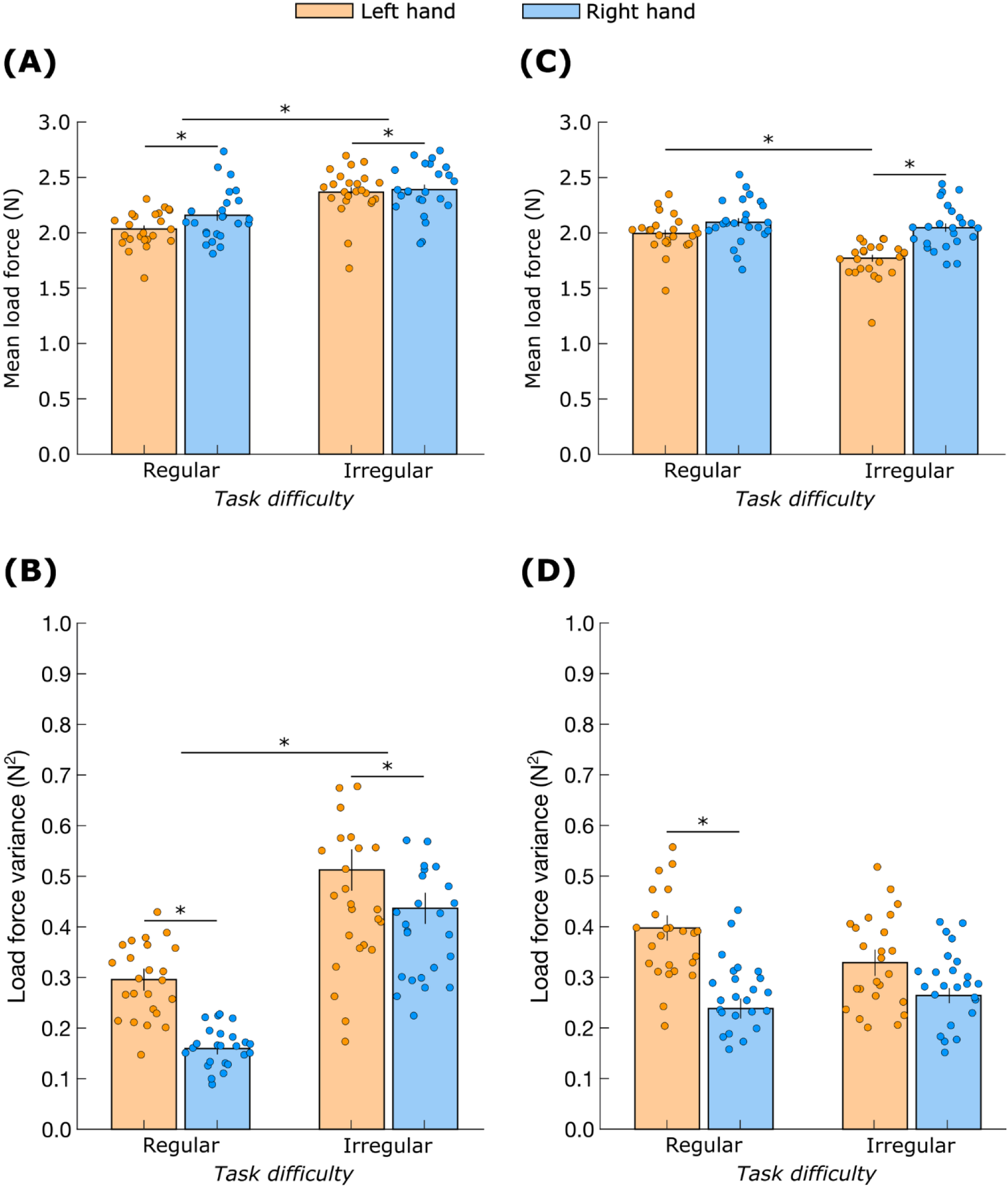
Load force characteristics. Mean (A) and variance (B) of the load on the moving object. Mean (C) and variance (D) of the load on the stabilized object. ‘*’ indicates significant differences (*p* < 0.05). Data are mean ± standard error.

The load force variance on the moving object (Fig 8B) was affected by *hand* (F_(1,23)_ = 19.0; *p* < 0.01), and *task difficulty* (F_(1,23)_ = 104.0; *p* < 0.01). Post-hoc analysis revealed that the load force variance was lower for the right object compared to the left (Cohen’s *d* = 0.9) and lower during regular movement compared to irregular movement (Cohen’s *d* = 2.1).

The mean load force on the stabilized object (Fig 8C) showed a *hand* × *task difficulty* interaction (F_(1,23)_ = 9.8; *p* < 0.01). Post-hoc analysis revealed that the mean load force was higher for the right object compared to the left but only during irregular movement (Cohen’s *d* = 1.4). Furthermore, the mean load force was higher during regular movement compared to irregular movement but only for the left object (Cohen’s *d* = 1.2).

The load force variance on the stabilized object (Fig 8D) showed a significant *hand × task difficulty* interaction (F_(1,23)_ = 6.4; *p* = 0.02). Post-hoc analysis revealed that the variance was higher for the left object compared to the right but only during the regular movement (Cohen’s *d* = 1.2).

### Task performance measures

The RMSE in the vertical direction for the moving object (Fig 9A) was not affected by *hand* (F_(1,23)_ = 0.01, *p* = 0.92) but was affected by *task difficulty* (F_(1,23)_ = 6128, *p* < 0.01). Post-hoc analysis revealed that the RMSE was low during regular movement compared to irregular movement (Cohen’s *d* = 16).

**Fig 9.**
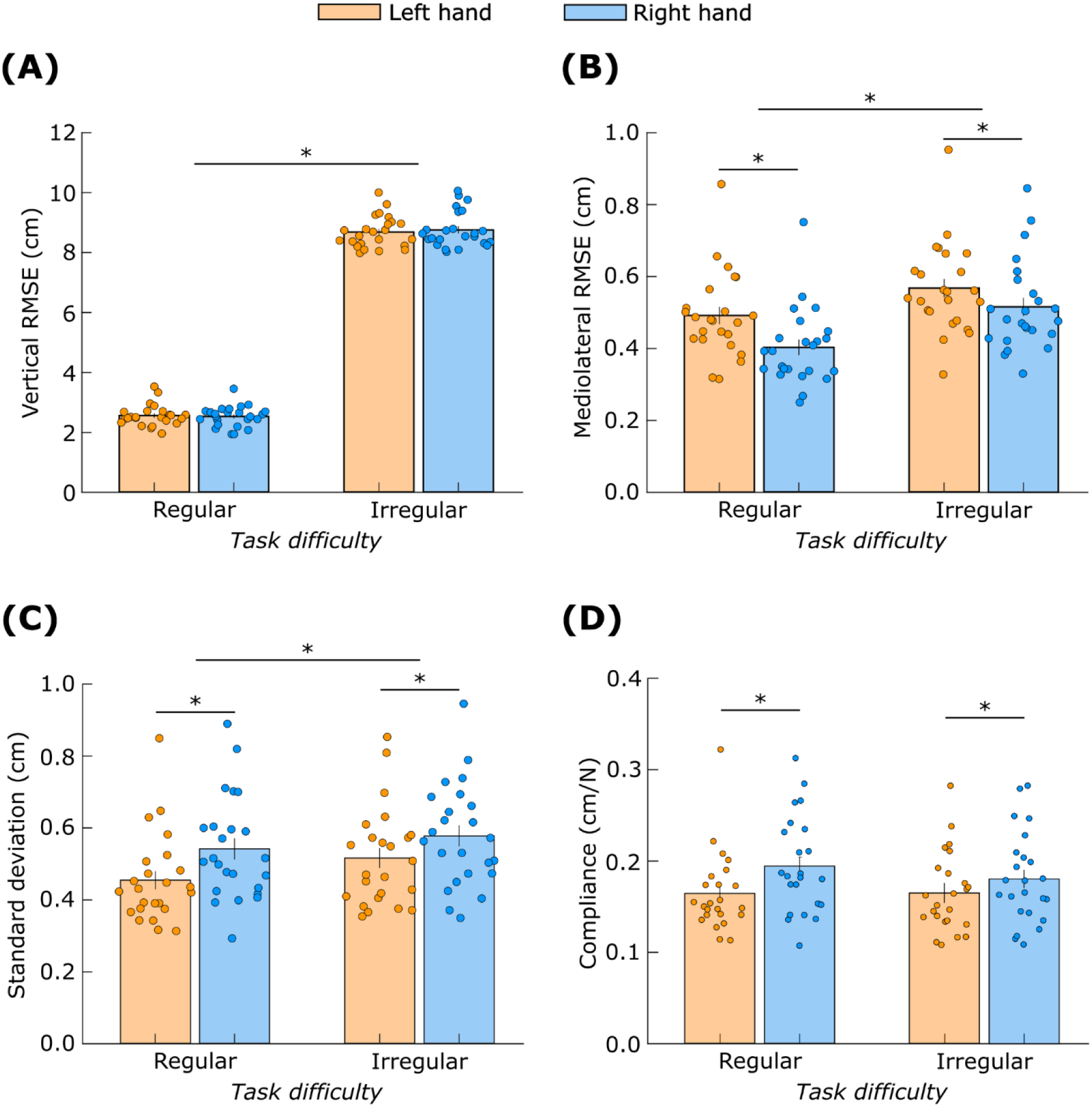
Task performance measures. RMSE (A) vertical and (B) mediolateral during object movement. (C) standard deviation and (D) compliance during object stabilization. ‘*’ indicates significant differences (*p* < 0.05). Data are mean ± standard error.

The RMSE in the mediolateral direction for the moving object (Fig 9B) was affected by *hand* (F_(1,23)_ = 30.1, *p* < 0.01) and *task difficulty* (F_(1,23)_ = 51.8, *p* < 0.01). The RMSE was low for the object in the right hand compared to the left hand (Cohen’s *d* = 1.1) and low for regular movement compared to irregular movement (Cohen’s *d* = 1.4).

The standard deviation of the stabilized object’s position (Fig 9C) was affected by *hand* (F_(1,23)_ = 18.0, *p* < 0.01) and *task difficulty* (F_(1,23)_ = 8.0, *p* < 0.01). Post-hoc analysis revealed that standard deviation was low for the object in the left hand compared to the right hand (Cohen’s *d* = 0.9) and low for regular movement compared to irregular movement (Cohen’s *d* = 0.6).

Finally, the compliance of the stabilized object’s position (Fig 9D) was affected by *hand* (F_(1,23)_ = 10.8, *p* < 0.01) but not by *task difficulty* (F_(1,23)_ = 0.9, *p* = 0.35). Post-hoc analysis revealed that the object stabilized by the left hand was less compliant than the object stabilized by the right hand (Cohen’s *d* = 0.7).

### Measures of grip force control

The grip-load correlation coefficient for the moving hand (Fig 10A) showed a significant *hand* × *task difficulty* interaction (F_(1,23)_ = 6.6, *p* = 0.02). Post-hoc analysis revealed that the coefficient was higher for the right hand compared to the left hand but only during the regular movement (Cohen’s *d* = 1.0). Furthermore, the coefficient was higher for the irregular movement compared to the regular movement but only for the left hand (Cohen’s *d* = 0.7).

**Fig 10.**
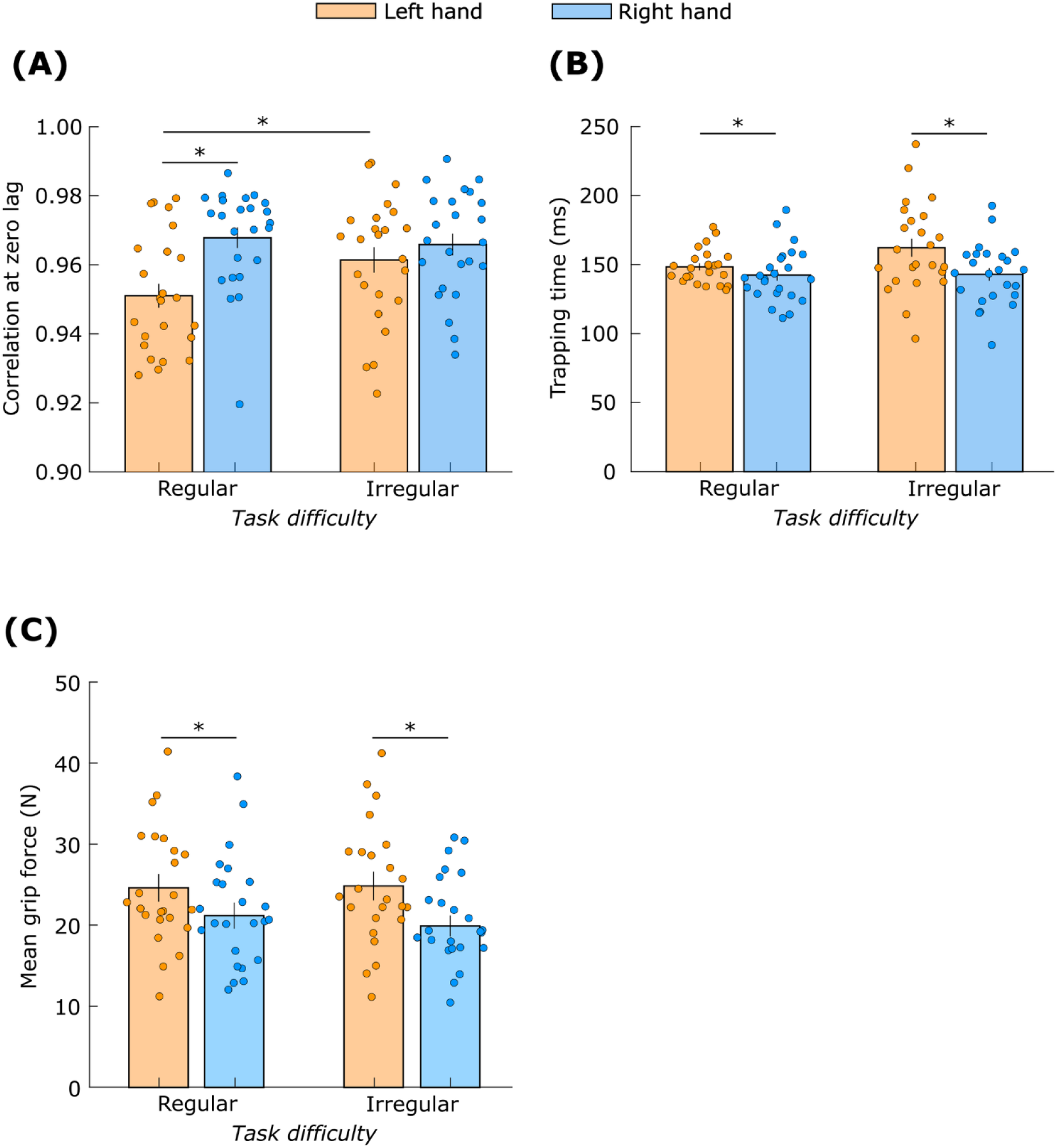
Grip force characteristics. Measures quantifying grip-load force coupling during object movement (A) Correlation at zero lag, and (B) trapping time. (C) Mean grip force during object stabilization. ‘*’ indicates significant differences (*p* < 0.05). Data are mean ± standard error.

The non-linear measure Trapping Time for the moving hand (Fig 10B) was affected by *hand* (F_(1,23)_ = 8.7, *p* < 0.01) but not by *task difficulty* (F_(1,23)_ = 1.7, *p* = 0.2). Post-hot analysis revealed that the trapping time was lower for the right hand than the left hand (Cohen’s *d* = 0.6).

The mean grip force on the stabilized object (Fig 10C) was affected by *hand* (F_(1,23)_ = 34.7, *p* < 0.01) but not by *task difficulty* (F_(1,23)_ = 0.55, *p* = 0.45). Mean grip force was higher for left hand compared to right hand (Cohen’s *d* = 1.2).

## Discussion

The primary aim of this study was to determine whether features of dynamic dominance extend to the movements of hand-held objects and to grip-force control. Our results support the predictions of the dynamic dominance theory. The right hand showed superior tracking performance in the mediolateral direction, indicating that the right hand produced straighter movements compared to the left hand, partially supporting hypothesis H1. However, there was no between-hand difference in the tracking performance in the vertical direction. The left hand showed superior stabilizing performance, supporting H2. The grip-load coupling strength during object movement was stronger in the right hand, supporting H3. Finally, the left hand exerted a higher grip force when stabilizing the object, supporting H4.

The secondary aim of this study was to determine whether task difficulty modulated dynamic dominance effects in grip-force control. Task difficulty had a minimal effect on grip force characteristics. The effect of task difficulty on grip-load coupling was not consistent, and the data provided weak support for H5. The linear correlation coefficient between grip and load forces increased with task difficulty but only for the left hand. The non-linear coupling measure, Trapping Time, was not influenced by task difficulty for either hand. Finally, task difficulty did not affect mean grip force when either hand stabilized the object; there was no support for H6.

### Evidence for dynamic dominance in grip force control

The brain’s hemispheric division of labor is an ancient feature predating vertebrates [62]. The left hemisphere handles well-established behaviors, while the right hemisphere detects and responds to unexpected stimuli [63-65]. The division enhances behavioral efficiency. These notions were extended to arm motion control when Sainburg and colleagues developed the dynamic dominance theory of handedness [8, 40]. They characterized the behavioral consequences of the lateralization in brain function by studying wrist trajectories in healthy individuals and stroke survivors [8, 12, 40, 66-70].

We wanted to determine whether hemispheric specialization was evident in the control of manipulated objects. The phenomenon of grip-load coupling suggests that the control of grip force is coupled to the control of arm motion, making it prima facie likely that dynamic dominance extends to grip force control. However, humans can alter grip-load coupling without changing arm posture. For example, grip force increases to maintain an object’s position in anticipation of an external disturbance [21] and also *before* voluntarily moving the arm to manipulate the object [22]. Furthermore, non-overlapping neural substrates are involved in the control of arm motion and digit forces. The reaching circuit in the cerebral cortex consists of the V6A, medial intraparietal area and dorsal premotor cortex [15-17], whereas the grasping circuit is comprised of the anterior intraparietal area, and ventral and dorsal premotor cortex [18, 20, 71, 72]. Therefore, the control of arm motion and grip forces is dissociated, and explicit behavioral evidence supporting dynamic dominance in grip force control was necessary. We provided this evidence, and it is our main contribution to the literature.

The stronger grip-load coupling in the right hand (Figs 10A and B) indicates that the left hemisphere’s specialization in predictive control influences grip forces. The between-hand difference in this coupling may be attributed to the differences in the efficacy of the predictive control. The existence of grip-load coupling implies that the reaching and grasping neural circuits interact [19]. Therefore, we speculate that the superior predictive control by the left hemisphere may arise from enhanced or better optimized interactions between the circuits in that hemisphere. Further research is necessary to test this hypothesis.

In addition to the stronger grip-load coupling, the object motion was straighter, i.e., with lower medial-lateral deviation, with the right compared to the left hand (Fig 9B). This result matches the straighter wrist paths while reaching with the right arm observed previously [8, 9]. The straighter movements suggest better prediction of the arm inter-segmental dynamics and the spring load by the left hemisphere, compatible with earlier work showing straighter paths of the right compared to the left wrist after adapting to predictable external force fields [40].

However, we did not observe between-hand differences in target tracking performance in the vertical direction (Fig 9A). This may be because target tracking requires controlling the object based on visual feedback on the target’s position, and hemispheric specialization for visual-spatial processing may have interfered with that for movement control. The right hemisphere is considered superior in visual-spatial processing [73], and it has been suggested that this specialization is evident when experiencing a mismatch between motor intention and visual feedback [74]. We speculate that in the right hemisphere, specialization of visual-feedback-driven tracking compensated for the weaker predictions of arm dynamics and spring loads. Whereas, in the left hemisphere, better predictions of the arm and spring dynamics compensated for the weaker visual-feedback-driven processes. This may have led to similar tracking performance by both hands.

The superior impedance control with the right hemisphere was manifested in higher grip force in the left hand when stabilizing the object. At the peripheral or mechanical level, higher grip force enhances mechanical impedance of the object because the object is less likely to slip from the fingers [28, 34], and because higher grip force involves greater co-contraction in the arm muscles, which, in turn, leads to higher endpoint impedance [29-31]. In particular, extrinsic hand muscles are involved in producing grip force with a pinch grasp [75, 76]. These muscles have their bellies in the forearm, with tendons crossing the wrist [77]. When these muscles contract to create a higher grip force, they also generate a larger wrist flexion moment. To maintain wrist orientation, the wrist extensors must contract, thus generating greater co-contraction. This stiffens the wrist and improves endpoint impedance [28, 29].

The higher grip force was accompanied by better stabilization (lower compliance) of the object in the left hand, indicating the efficacy of the right hemisphere at impedance control. This suggests better manipulation of long-latency reflexes, mediated by motor cortical pathways [78, 79], by the right hemisphere. Higher gain of the stretch reflex implies heightened sensitivity of the spinal motor neurons [80, 81], which would enhance muscular responses triggered by proprioceptive feedback [28, 82-84], eventually resulting in stronger resistance to external perturbations [85, 86] by the left hand.

### Effect of task difficulty on bimanual grip force control

Contrary to our expectation, *task difficulty* had a minimal effect on between-hand differences in grip force characteristics. The hypothesized increase in the grip-load coupling for the irregular motion compared to the regular motion was not consistent, lending only partial support to earlier findings [33]. One reason could be that there was no external load on the object in the previous work [33, 87, 88], whereas the spring that coupled the two objects induced an external load on the moving object. Furthermore, the load force in the irregular task was less predictable, which may have led to weaker grip-load coupling than expected. We verified that the excursions of the stabilized object that occurred despite the participants’ efforts were more complex for the irregular task (supplemental material 1). Furthermore, the total load on the moving object was dominated by the spring load: the average spring force (2.95 N) was 17 times larger than the average inertial force (0.17 N), and the variability in the spring load (0.66 N^2^) was 33 times larger than that in the inertial load (0.02 N^2^) on the moving object (supplemental material 2). Therefore, greater irregularity (or complexity) in the excursions of the stabilized object likely influenced the predictability of the spring and total load on the moving object, which may have negatively impacted the grip-load coupling for the irregular task.

The mean grip force on the stabilized object did not increase during the irregular task compared to the regular task. This result contradicts the finding from a recent study that the mean grip force is three times more sensitive to variability rather than the average value of environmental perturbations [34]. A likely reason could be that the spring load on the stabilized object was not sufficiently unpredictable for the grip force to show significant increase across tasks. The load on the stabilized object was due to voluntary movement of the moving hand exerted via the spring. Furthermore, the spring stiffness was not modulated across trials. Therefore, the motor system could have predicted the spring load arising from the arm motions even during the irregular task. Thus, it is plausible that both feedforward and feedback mechanisms were involved when stabilizing the object against perturbations arising from volitional actions [89], and therefore the effect of load force variability on mean grip force was not observed.

Our plausible explanation for the lack of effect on grip-load coupling is too much unpredictability in the load on the moving object, whereas for the grip force, it is too much predictability in the load on the stabilized object. This might appear incompatible. However, the unpredictable component of the load on the moving object is thought to arise from *involuntary* movements of the stabilized object, likely due to motor noise and mechanical disturbances in the apparatus. In contrast, the loads on the stabilized object are thought to arise mainly from the *voluntary* movement of the other arm, which the nervous system can predict.

### Contributions to Grip Force from the Central Drive

Although our data demonstrates hand-specific differences in object manipulation and grip force control that are consistent with the dynamic dominance theory, inter-hemispheric interference/coupling should also be considered while explaining our results [41, 42, 90, 91]. When two hands simultaneously perform asymmetrical tasks, the grip force of both hands tends to be more similar than expected. For example, in a bimanual task in which each hand moves an object of the same weight in one condition, increasing the weight of just one object leads to an increase in the grip force of both hands. This interference occurs because of a common central drive, hypothesized to facilitate bimanual control, that increases the grip force in both hands [41]. It is possible that the grip forces of both hands in our study are affected by the common drive, and therefore, the hand-specific differences in grip force characteristics may not be reflection of just hemispheric specialization.

Nevertheless, hemispheric specialization likely contributes to the patterns in our data. The central drive is stronger and leads to greater equalization of grip forces when the two hands perform asymmetrical tasks, for example, when one hand moves one object while the other hand holds another object static [91], or when both hands move two objects simultaneously but with different loads [41, 42]. However, the effects of the central drive are the same when the roles of the hands are switched. [41, 42]. In contrast, the mean grip force in our study is different when the left versus the right hand stabilized the object (Fig 10C) indicating that both dynamic dominance and the common central drive influence grip force control. Future work should quantify the contributions of these two phenomena and their interactions in ecological bimanual tasks.

### Left-handed individuals

A clear limitation of this study is the exclusion of left-handed individuals. Left-handed individuals show more diverse behavior and neurophysiology compared to their right-handed counterparts [92, 93]. The laterality indices for left-handed individuals are more variable [94], and left handers demonstrated more balanced hand use indicating a lack of bias toward one hand [93]. This pattern is likely related to left handers exhibiting higher activation of the ipsilateral cortex during unimanual tasks compared to right handers [92]. This suggests that the more symmetrical behavior in left-handers may result from more bilateral hemisphere recruitment, allowing each hand to benefit from specialized functions of each hemisphere [95]. Future research should investigate neural mechanisms of grip force control in left-handed individuals.

## Conclusion

This study demonstrates that the grip fore control and object manipulation during bimanual tasks reflect dynamic dominance. The object movement was straighter, and the grip-load coupling was stronger in the right hand, indicating superior predictive control of the right hand by the left brain hemisphere. In contrast, the object compliance was lower, and the mean grip force on the object was higher in the left hand, indicating superior impedance control of the left hand by the right brain hemisphere. Task difficulty had a weak effect on grip-load coupling during object movement and no effect on mean grip force during object stabilization, likely due to task design where the objects in the two hands were coupled with a spring. The load on the moving object and thereby the grip-load coupling was impacted by the complexity of inadvertent excursions of the stabilized object, and external load on the stabilized object was only partially unpredictable since it arose from the voluntary movement of the other hand. Overall, this study highlights hand-specific specialization of the control of prehensile manipulation and extends the domain of applicability of the dynamic dominance theory to bimanual prehension.

