## Supplementary material for "Handy divisions: Hand-specific specialization of prehensile control in bimanual tasks": Supplimental material

### Supplementary materials

#### 1. Complexity of the excursions of the stabilized object

The stabilized object displayed some excursions. We assessed whether these excursions were more complex in one of the tasks. We computed sample entropy of the displacement of the stabilized object in the mediolateral and the vertical directions as the measure of complexity. The time series was divided into vectors of length $m$ (set to m=3), and each vector was compared with subsequent vectors for repeating elements within a specified tolerance $r$ (set to $r = 2$) [96]. Such subsequent comparisons are repeated with vectors of length $m$+1. Sample entropy was then computed as the negative natural logarithm of the conditional probabilities of these two comparisons. A higher sample entropy implies higher complexity.

We pooled sample entropy values across hands for the regular and the irregular tasks. Two-sample t tests revealed that the sample entropy was higher ($t_{(23)} = 11.1$; $p < 0.01$) during irregular (0.13±0.01) than the regular motion (0.09±0.01) in the mediolateral direction. Likewise, the sample entropy was higher ($t_{(23)} = 12.5$; $p < 0.01$) during irregular (0.12±0.01) than irregular motion (0.08±0.01) in the vertical direction.

Therefore, the excursions of the stabilized object were more complex during the irregular task.

#### 2. Comparisons of load force components for the moving hand

Here, we report the mean and variance of components of load force – inertial and spring forces – for each hand during object movement. Linear mixed-effects models were fitted with three experimental factors (*hand* × *task difficulty* × *load component)* and their two- and three-way interactions as fixed effects and participants are a random effect. Tukey-Kramer test was used to perform post-hoc pairwise comparisons when significant interaction effects were observed. Effect sizes were quantified by computing Cohen's *d*. Significance was set at an α-level of 0.05.

The mean force (Fig S1A) showed a significant *hand* × *load component* interaction ($F_{(1,23)} = 4.92$, $p = 0.03$). Post-hoc analysis revealed that only the mean inertial force was higher in the right hand than the left hand (Cohen's $d$ =0.5). The mean force also showed a significant *task difficulty* × *load component* interaction ($F_{(1,23)} = 223.0$, $p < 0.01$). Post-hoc analysis revealed that the mean spring force was greater than the mean inertial force. However, the difference between the two mean forces was higher during irregular movement (Cohen's $d = 2.2$).

The force variance (Fig S1B) showed a significant *hand* × *load component* interaction ($F_{(1,23)} = 11.3$, $p < 0.01$). Post-hoc analysis revealed that only the inertial force variance was higher in the right hand than the left hand (Cohen's $d$ =0.6). The force variance also showed a significant *task difficulty* × *load component* interaction ($F_{(1,23)} = 77.7$, $p < 0.01$). Post-hoc analysis revealed that the spring force variance was greater than the inertial force variance. However, the difference between the two force variances was higher during irregular movement (Cohen's $d = 3.3$).

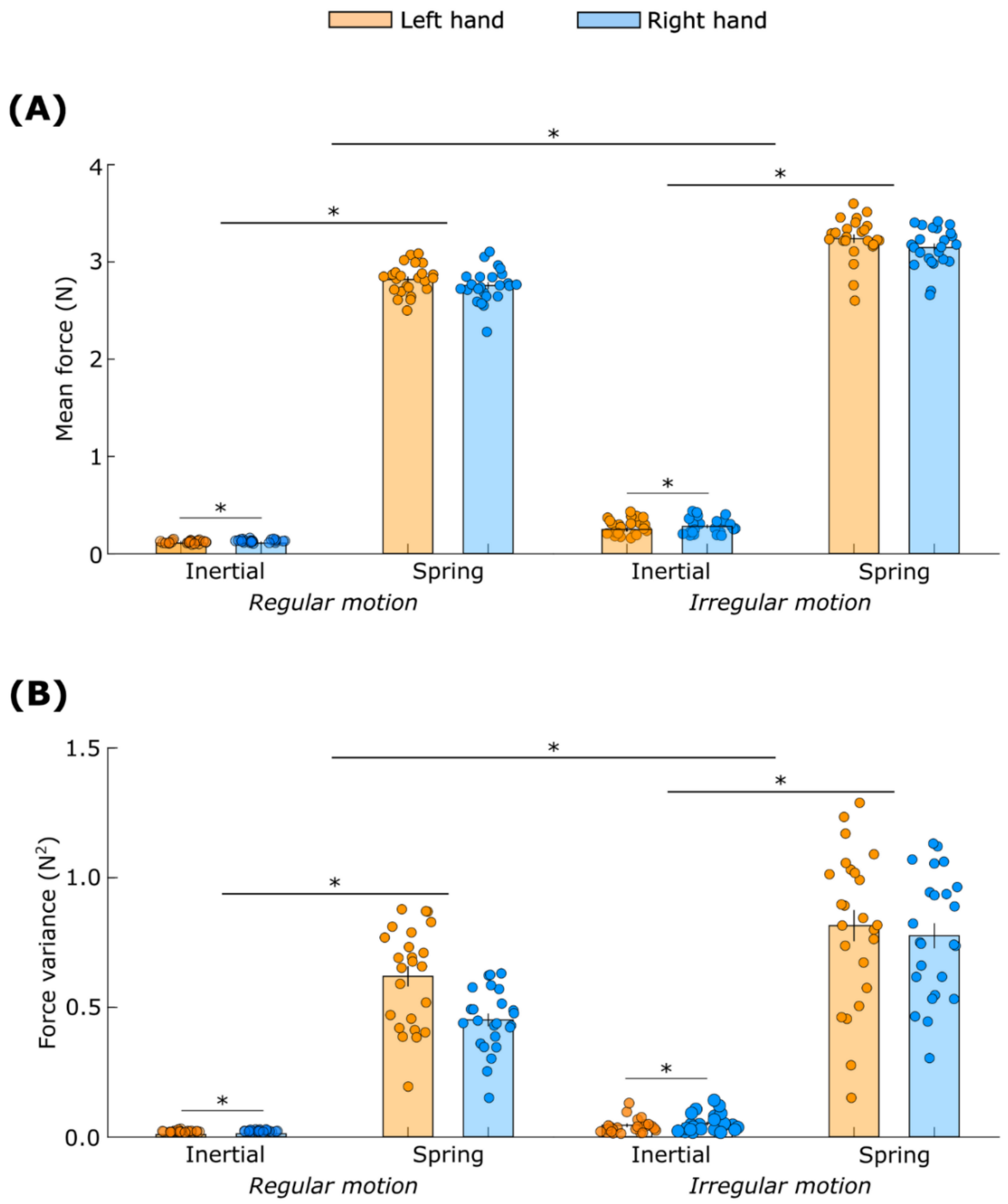


**Fig S1. Load force components**. (A) mean and (B) variance of forces. '*' indicates significant differences ($p < 0.05$). Data are mean ± standard error.

#### 3. Performance scores

Participants repeated the same movements 10 times in various tasks. Therefore, to check whether participants were able to predict the target's position in each *task type*, a learning analysis was performed for each *hand* × *task type* block. To test whether learning occurred, a simple linear regression analysis was performed within each *hand* × *task type* block, with trial number as the independent variable and the score as the dependent variable. A significant negative slope would indicate learning occurred after repeated exposure to the target profile. The results are summarized in Table A.2.

In the *left-hand* × *irregular task* block, the performance score improved in six participants. In the *right-hand* × *irregular task* block, the performance score improved in six participants. In the *left-hand* × *regular task* block, the performance score improved in four participants. In the *right-hand* × *regular task* block, the performance score improved in eight participants. Two participants demonstrated learning effects in three of the four tasks and four participants demonstrated learning effects in two of the four tasks.

Overall, only a minority of participants demonstrated learning in each of the tasks.

**Table S1. Summary statistics for the learning effect across trials in each *hand* × *task type* block.** A $p$-value < 0.05 indicates slope that is significantly different than zero. A significant negative slope indicates an effect of learning and is denoted by *****.

| | *Left hand moving × irregular task* | | *Right hand moving × irregular task* | | *Left hand moving × regular task* | | *Right hand moving × regular task* | |
|---|---|---|---|---|---|---|---|---|
| Participant # | Slope ($cm^2$/trial) | $p$-value | Slope ($cm^2$/trial) | $p$-value | Slope ($cm^2$/trial) | $p$-value | Slope ($cm^2$/trial) | $p$-value |
| 1 | 0.42 | 0.46 | 1.68 | 0.16 | 0.17 | 0.33 | 0.33 | <0.01 |
| 2 | -2.29* | 0.01 | -0.28 | 0.84 | 0.00 | 1.00 | -0.53* | 0.01 |
| 3 | -0.40 | 0.65 | -0.75 | 0.40 | 0.05 | 0.74 | -0.30 | 0.07 |
| 4 | -0.04 | 0.89 | 0.40 | 0.49 | -0.18 | 0.35 | 0.00 | 0.98 |
| 5 | 0.02 | 0.97 | -0.72 | 0.28 | -0.21 | 0.34 | -0.16 | 0.53 |
| 6 | -2.31 | 0.16 | -3.83 | 0.05 | -0.79* | 0.04 | -0.71 | 0.05 |
| 7 | -0.89 | 0.32 | -1.29 | 0.21 | 0.06 | 0.76 | -0.40 | 0.10 |
| 8 | -3.74* | 0.04 | -3.21* | <0.01 | -1.01* | <0.01 | -0.58 | 0.05 |
| 9 | -3.77* | 0.01 | -0.79 | 0.46 | -0.74* | <0.01 | -0.11 | 0.70 |
| 10 | -0.67 | 0.43 | -0.78 | 0.26 | -0.64 | 0.08 | 0.41 | 0.20 |
| 11 | -1.46 | 0.06 | -1.54 | 0.06 | -0.38 | 0.13 | -0.46 | 0.08 |
| 12 | -2.59 | 0.11 | -3.03 | 0.06 | -0.50* | <0.01 | -0.05 | 0.54 |
| 13 | -1.45* | 0.01 | -1.08* | <0.01 | -0.17 | 0.38 | -0.79* | <0.01 |
| 14 | 0.34 | 0.74 | -2.05* | 0.01 | -0.27 | 0.16 | -0.40 | 0.10 |
| 15 | -0.63 | 0.67 | -5.81* | <0.01 | -0.40 | 0.26 | -0.15 | 0.42 |
| 16 | -0.25 | 0.68 | -1.05 | 0.07 | -0.40 | 0.36 | -0.18 | 0.20 |
| 17 | -0.05 | 0.83 | -1.87* | <0.01 | -0.02 | 0.91 | -0.70* | <0.01 |
| 18 | -1.15* | 0.02 | -0.71 | 0.54 | 0.00 | 0.99 | -0.34 | 0.33 |
| 19 | -1.46 | 0.08 | -0.83 | 0.19 | -0.18 | 0.14 | -0.65* | <0.01 |
| 20 | -0.91 | 0.33 | 0.93 | 0.69 | -0.18 | 0.12 | -0.34 | 0.06 |
| 21 | -1.56 | 0.24 | -0.19 | 0.89 | 0.28 | 0.44 | -0.51* | 0.01 |
| 22 | -2.75* | 0.04 | -2.78* | 0.03* | -0.31 | 0.43 | -0.16 | 0.43 |
| 23 | -0.22 | 0.69 | -1.33 | 0.05 | -0.24 | 0.15 | -0.70* | 0.02 |
| 24 | -2.00 | 0.07 | -1.18 | 0.23 | -0.27 | 0.17 | 0.03 | 0.89 |
